# BMP and NODAL Paracrine signalling regulate the totipotent-like cell state in embryonic stem cells

**DOI:** 10.1101/2025.10.14.682333

**Authors:** Sanidhya Jagdish, Loick Joumier, Sabin Dhakal, Gilberto Duran-Bishop, Mohammed Usama, Mohan Malleshaiah

**Affiliations:** Institut de recherches cliniques de Montréal (IRCM); Department of Experimental Medicine, McGill University; Département de biochimie et médecine moléculaire, University of Montréal

**Author notes:** Equal contribution (Sanidhya Jagdish, Loick Joumier).

## Abstract

Cell–cell communication coordinates signalling between cells to guide context-dependent cell fate decisions such as proliferation, differentiation, and lineage specification. Such communication mechanisms are poorly understood in regulating the stem cell states. In this study, we investigate how cell-cell communication regulates cell fate transitions in heterogeneous embryonic stem cell populations, with a particular focus on totipotent-like cells that resemble the two-cell stage embryo. Using single-cell RNA sequencing in combination with computational frameworks, we map ligand– receptor interactions and model downstream regulatory effects across various stem cell states. We functionally validate the predictions by selectively perturbing signalling pathways under specific culture conditions. Our data reveal the key roles of BMP and NODAL (TGF-β) signalling in mediating intercellular communication to shape stem cell identity and heterogeneity. These findings enhance our understanding of the signalling logic that governs early developmental cell fate decisions, providing new insights into stem cell biology with broad implications for regenerative medicine and developmental modelling.

## 2) INTRODUCTION

Cell fate determination is fundamental to embryonic development, where a single cell gives rise to all specialized lineages of an organism (1). This process is governed by intricate regulatory networks that integrate transcriptional, epigenetic, and environmental signals (2). The accurate coordination of these developmental pathways is essential not only for normal embryonic development but also for maintaining physiological balance (homeostasis) and supporting tissue regeneration in adulthood (3). Cells act as dynamic systems, with complex signalling pathways that are closely interconnected through feedback loops and crosstalk (3). Systems biology approaches involving single-cell measurements are thus crucial for unravelling the sophisticated mechanisms underlying cell fate decisions (3, 4).

In multicellular systems, cell identity and function are controlled by signals from neighbouring cells, creating a dynamic microenvironment (5). Such cell–cell communication (CCC) is facilitated by secreted proteins, like growth factors and cytokines, which activate signalling cascades in adjacent cells, leading to changes in their gene expression, epigenetics, metabolism, and functional patterns (6-8). CCC occurs in several modes: autocrine (self-regulation), paracrine (localized responses), endocrine (long-range hormonal signalling), and synaptic (rapid neural communication) (9). At the population level, CCC can coordinate collective behaviours, such as morphogenesis, wound healing, and immune responses (6), while also producing heterogeneity through signalling gradients (10). Therefore, a detailed understanding of intercellular communication mechanisms is essential for fundamental biology and therapeutic development.

The CCC mechanisms are particularly evident in stem cell niches, where local signalling controls self-renewal, lineage commitment, and plasticity (11, 12). Due to their ability to differentiate into specialized cell types, stem cells provide a suitable model for studying CCC mechanisms in developmental contexts. Pluripotent cells, such as embryonic stem cells (ESCs) and induced pluripotent stem cells (iPSCs), represent the stem cell niche of early embryogenesis (13, 14). Their ability to self-renew and differentiate into all embryonic cell types provides a proxy for examining how CCC mechanisms regulate the emergence of lineage-specific cell types (15). The establishment and maintenance of the pluripotent stem cell state itself is regulated by CCC. For instance, fibroblast growth factor (FGF) signalling controls the balance between self-renewal and differentiation of ESCs: activation of the FGF–ERK pathway promotes differentiation, while its inhibition preserves pluripotency (16). Similarly, paracrine signalling through bone morphogenic protein (BMP) also regulates the pluripotent and other heterogeneous cell states of ESCs (17). ESCs are inherently heterogeneous and often display distinct cell states of early embryogenesis (17, 18): for example, totipotent-like, naïve pluripotent, primed, and primitive endoderm (PrEn) cell states. Such heterogeneity impacts the reproducibility of ESC-based models and therapies (19), but also mirrors the lineage diversity in the early mammalian embryo, making ESCs a physiologically relevant system for studying how distinct cell populations communicate and influence one another (20). Thus, ESC heterogeneity provides a unique opportunity to dissect CCC within a controlled, developmentally relevant context.

Of particular interest in the ESC system are totipotent-like cells (TLCs), which display transcriptional signatures reminiscent of the two-cell (2C) stage embryo and can contribute to both embryonic and extraembryonic lineages (21). Although rare and transient, their emergence highlights the plasticity of ESC cultures and the influence of extrinsic signals on their identity. Despite the importance of totipotent cells to initiate mammalian embryonic development and give rise to both embryonic and extraembryonic cell types, the regulatory mechanisms controlling the totipotent cell state are poorly understood. Understanding how TLCs maintain or exit their state is thus key to elucidating early embryo development.

Studying CCC among specific cell states requires high-resolution single-cell measurements. Single-cell RNA sequencing (scRNA-seq) enables detailed profiling of cell states, including their signalling and transcriptional responses (22). To estimate the extent of specific ligand-receptor interactions among cell states from scRNA-seq data, computational frameworks such as CellChat (23) and NicheNet (24) have been developed. Furthermore, the influence of paracrine signalling on the transcriptional regulatory networks that determine cell fate can be modelled using methods like CellOracle (25). Thus, the effective use of scRNA-seq in combination with computation can enable the systematic mapping of ligand–receptor interactions, modelling of signalling and transcriptional networks, and prediction of functional outcomes, offering an integrated systems-level approach to understanding how extrinsic signals are processed at the single-cell level in cell fate determination. In this study, we used scRNA-seq data, coupled with computational frameworks such as CellChat (23), NicheNet (24), and CellOracle (25), to infer CCC among cell states of mouse ESCs. We mapped the paracrine signalling landscape of TLCs and identified BMP and NODAL as the key CCC routes to either enhance or diminish the TLC state, respectively.

## 3) MATERIALS AND METHODS

### Single-Cell RNA-Seq Analysis

scRNA-seq data were processed and analyzed in R (v4.5.0) using the same pipeline described in our previous study (17). All preprocessing, normalization, and statistical thresholds were applied identically to ensure consistency across datasets.

### Cell–Cell Communication Analysis

Intercellular communication was investigated using CellChat, NicheNetR, and CellOracle, each providing complementary insights into ligand–receptor signalling and gene regulatory dynamics.

### CellChat Analysis

CellChat (v1.6.1) infers communication networks by modelling interactions between sender and receiver cell populations using a curated database of ligand–receptor pairs (24). This framework allows for the identification of dominant signalling pathways and quantification of their contribution to overall intercellular communication. We applied CellChat to quantify interaction networks, determine pathway activity, and compare global signalling patterns between LB and LBPXRS conditions.

### Differential NicheNetR Analysis

NicheNetR (v2.2.1) was used to link ligand activity in sender cells with downstream transcriptional responses in receiver cells (25). For differential analysis, we defined four distinct signalling niches:

1. *Totipotent niche* – totipotent cells as receivers; primed, PrEn, and pluripotent cells as senders.
2. *Pluripotent niche* – pluripotent cells as receivers; primed, PrEn, and totipotent cells as senders.
3. *Primed niche* – primed cells as receivers; totipotent, PrEn, and pluripotent cells as senders.
4. *PrEn niche* – PrEn cells as receivers; primed, pluripotent, and totipotent cells as senders.

Differential ligand–receptor analysis was performed with a log fold-change cutoff of 0.15. The top 250 predicted target genes were retained for prioritization. Ligand–receptor pairs were ranked according to recommended DE-NicheNetR thresholds, and the top 25 ligands were displayed for visualization.

### Pathway-Focused NicheNetR Analysis

We next performed a pathway-specific NicheNetR analysis focusing on signalling cascades relevant to early development, including BMP, FGF, WNT, TGF-β, and PDGF pathways (17). The prioritized ligand-receptor pairs were selected from each pathway and merged into one matrix. Ligand–receptor pairs were then filtered and prioritized based on expression levels and predicted activity using an interaction strength threshold of 0.4. We then generated curated dot plots to map the expression of selected ligands and receptors across sender and receiver populations in the LB condition. Pairs were retained if detected in TLC-to-cell communication with at least two of three receiver populations and were exclusive to either the TLC sender or receiver groups.

### Validation Experiments

R1 line of ESCs was cultured in LB media (N2B27 medium supplemented with LIF and BMP4). To assess the contribution of individual signalling pathways, cells were treated with the following inhibitors for 48 hours: LY2109761-0.8uM (TGFBRI/II inhibitor).

Optimal concentrations for each compound were determined by dose–response assays, with western blot analysis used to evaluate downstream pathway activity relative to control (LB condition). The most suitable treatment concentrations were selected based on these results. Subsequently, we examined the impact of pathway inhibition on TLC percentage by flow cytometry and evaluated transcriptional changes in cell state markers and downstream pathway targets by qPCR.

For protein extraction, cells were lysed in a 2× Laemmli buffer. Total RNA was isolated using TRIzol reagent and used for qPCR analysis. The percentage of MERVL-tdTomato-positive cells was quantified using flow cytometry to determine the proportion of TLCs under the indicated experimental conditions.

### Pseudotime inference

Trajectories were constructed using the Monocle 3 framework (v1.3.7) (26). scRNA-seq matrices were imported from the analysis performed in our previous *Cell Reports* paper (17). The root of the trajectory was defined as the terminal branchpoint nearest to the totipotent-like population, ensuring that the pseudotime variable reflected cell state transitions along natural embryonic development.

### In silico gene perturbation

In silico gene perturbation analysis was performed with CellOracle (v0.18.0) (25). Cell state-specific Gene Regulatory Networks (GRNs) were first reconstructed after loading CellOracle’s pre-built base GRN using the “load_mouse_scATAC_atlas_base_GRN” function. The pseudotime variable from the Monocle 3 analysis was implemented in the CellOracle object. Perturbed vector fields were generated for transcription factors, cell state markers, or targets of signaling pathways. Knockouts were simulated using the “simulate_shift” function, setting the target gene expression to 0. Grid parameters were set to 40 grid points and a cell density of 0.01 for digitizing the vector field.

## 4) RESULTS

### Extensive cell–cell communication supports ESC heterogeneity

Intercellular communication is mainly mediated by the interaction of ligands secreted or produced by the “sender” cells with specific receptors on the “receiver” cells. Such ligand-receptor interaction leads to activation of specific cell signalling pathways and their downstream effects in receiver cells. Since the pluripotent cell state in ESCs is sensitive to paracrine signalling effects, we hypothesized that the TLC state is similarly governed by its CCC with other cell states of ESCs (i.e. pluripotent, primed and PrEn) **(Fig. 1)**. We envisioned that the extent of the TLC state in the system is regulated by the extent of the sender-receiver communication between TLCs and other cell states via the ligand-receptor interactions.

**Figure 1:**
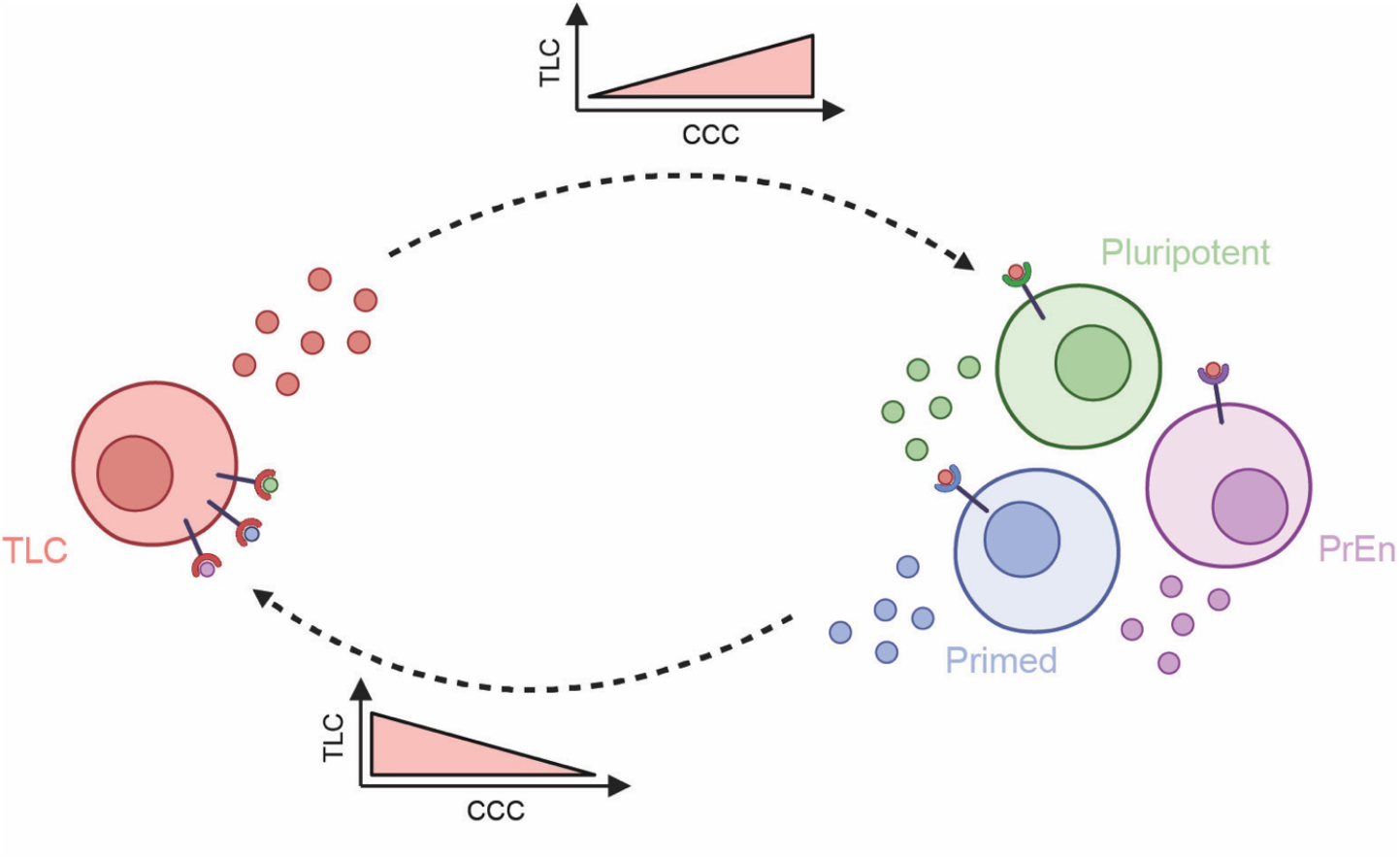
Schematic for the proposed model of signalling interactions regulating the TLC state and its exit. Totipotent-like cells (TLCs) secrete specific ligands that act in a paracrine manner by engaging receptors on neighbouring cell types, thereby reinforcing the TLC state and contributing to a supportive local signalling niche. In contrast, ligands predominantly secreted by pluripotent, primed, or primitive endoderm (PrEn) cells interact with receptors on TLCs to trigger their exit from the TLC state. Arrows indicate the directionality of signalling interactions.

Mouse ESCs are regulated by multiple cell signalling pathways (27). To understand the full extent of CCC mediated by signalling pathways, we first evaluated the previously reported scRNA-seq data of ESCs cultured in LIF and BMP growth factors (henceforth referred to as LB) (17). As expected, we observed all four cell states —TLCs, pluripotent, primed, and PrEn —in ESCs cultured in LB (17) **(Fig. 2A)**. This distribution indicates that LB supports broad state diversity and maintains cellular heterogeneity. Consistent with this, intercellular communication analysis using CellChat revealed a highly interconnected signalling network in the LB condition **(Fig. 2B)**. Each cell state received and transmitted paracrine signals, confirming extensive interactions across the system. The number of edges between nodes (e.g., a total of 30 signals between primed to PrEn, 15 signals between TLCs to PrEn, and 25 signals between pluripotent to PrEn) indicates that PrEn cells are major hubs of communication, receiving inputs from and sending outputs to all other states. We further observed bidirectional communication between TLCs and all other cell states, suggesting that their maintenance relies on multiple regulatory interactions, with a preference for PrEn.

**Figure 2:**
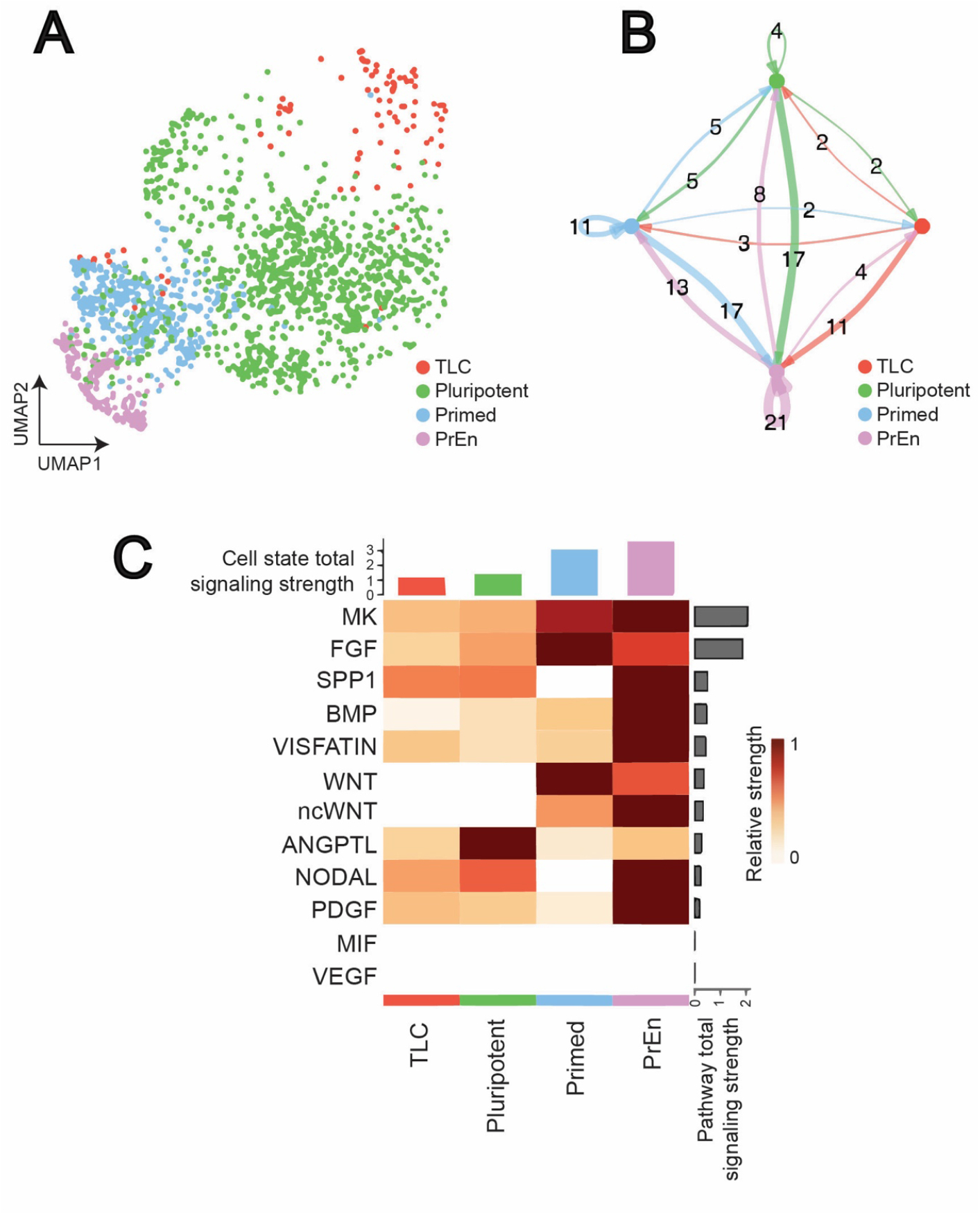
Intercellular communication landscape of ESCs in LB condition revealed by scRNA seq. **(A)** UMAP visualization of single-cell RNA-seq data of ESCs comparing the cellular landscape under the LB condition, highlighting major cell populations. **(B)** Circle plots depicting the number of inferred intercellular interactions among cell types in the LB condition as computed by CellChat. Edge thickness corresponds to interaction strength, while the arrow signifies the direction of the communication. **(C)** Overview of signalling pathway involvement in LB. The bar plot at the top shows the total signalling strength of a cell group by summarizing all signalling pathways displayed in the heatmap. The right grey bar plot shows the total signalling strength of a signalling pathway by summarizing all cell groups displayed in the heatmap.

A closer examination of signalling patterns further highlighted the complexity of this communication. Estimating the total signalling strength of each cell state and heatmap visualization of top contributing pathways revealed that primed and PrEn cells contributed the majority of signalling output **(Fig. 2C)**. Pathways such as MK, FGF, SPP1 and BMP were some of the dominant drivers of communication, primarily originating from the primed and PrEn cells. We decided to focus on FGF, BMP, NODAL (TGF-β), WNT, and PDGF as these have been established to regulate ESCs (17).

To specifically assess how TLCs are regulated within this environment, we performed receiver niche analysis for each cell state using NicheNet (24). NicheNet is a computational method that predicts how ligands from one cell type influence gene expression in another cell type. It integrates ligand– target regulatory networks with transcriptomic data to study cell–cell communication and its downstream effects. TLC-receiver analysis identified preferentially enriched ligands in pluripotent, primed and PrEn cells, and their corresponding receptors in TLCs **(Fig. 3A)**. Key ligand-receptor pairs included *Bmp4–Acvr2b, Wnt4–Lrp5, Tgfβ2–Acvr1b*, and *Lefty2–Acvr2b*, many of which belong to the FGF, WNT, BMP, and NODAL (TGF-β) families (28-30). These ligands were expressed at higher levels in sender populations and predicted to activate their corresponding pathways in TLCs. *Bmp2/4, Tgfβ2*, and *Lefty2* all belong to the TGF-β superfamily, which is crucial for stem cell maintenance, suggesting their role in regulating the TLC state (17, 31, 32). This data also revealed that PrEn and primed cells provided the strongest input ligand signals to influence TLCs. In addition to these canonical pathways, we also observed adhesion-related ligands (e.g., *Ccn1, Flrt3*), suggesting that cell–cell contact cues may further modulate TLC plasticity (33, 34). Similar analysis for pluripotent, primed and PrEn as receiver cells revealed the extensive ligand-receptor pairs of FGF, BMP, NODAL (TGF-β), WNT, and PDGF pathways involved in regulating the stem cell states **(Fig. 3B-D)**.

**Figure 3:**
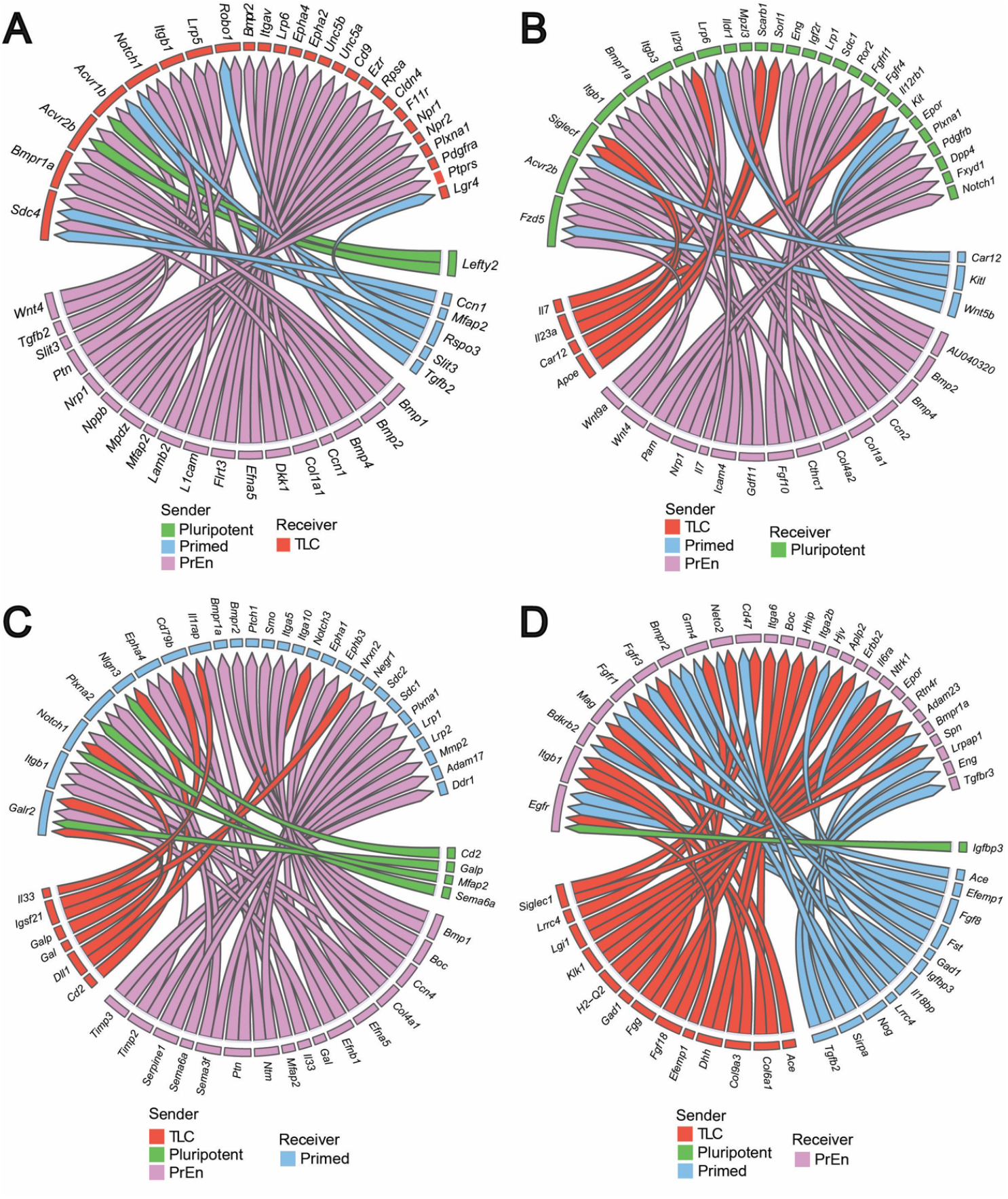
Differential NicheNet analysis reveals the top ligand-receptor pairs in the LB condition. Circos plots illustrating ligand–receptor interactions between all sender cell types and the **(A)** TLC, **(B)** pluripotent, **(C)** primed, and **(D)** PrEn receiver population in LB. Chords represent the strength and directionality of signalling inputs.

Taken together, these data reveal that cell states within ESC populations communicate via an intricate network of paracrine signalling. Higher level of CCC correlates with high cell state heterogeneity in ESCs, suggesting a regulatory relationship. They also show that the extent of the TLC state is likely heavily influenced by paracrine signalling from pluripotent, primed, and PrEn cells.

### Diminished CCC correlates with reduced heterogeneity

To validate whether the extent of signalling activity observed in ESC culture regulates cell state heterogeneity, we next evaluated the previously reported scRNA-seq data of ESCs cultured in LIF and BMP growth factors, as well as PD0325901, XAV939, Resorcyclic lactone, and SB431542 small molecules (henceforth referred to as LBPXRS) (17). In this condition, BMP-mediated cross-activation of the FGF, WNT, and NODAL (TGF-β) pathways is selectively inhibited using PD0325901, XAV939, and Resorcyclic lactone/SB431542, respectively (17). The LBPXRS condition has also been demonstrated to strikingly alter cell state composition by enriching TLCs and pluripotent cells and nearly eliminating primed and PrEn cells (17) **(Fig. 4A)**. Quantitative assessment of cell-type proportions confirmed these observations **(Fig. 4B)**: compared to LB, LBPXRS culture showed a ∼23% enrichment of TLCs and a ∼9% increase in pluripotent cells, accompanied by a drastic reduction of primed cells to 0.7% and PrEn to 0.3% of the population. These findings indicate that reduced signalling in the LBPXRS condition narrows the cell identity landscape and diminishes cellular heterogeneity. Thus, with overall low signalling activity and altered cell state composition, scRNA-seq data from LBPXRS is highly suitable for evaluating the role of CCC.

**Figure 4:**
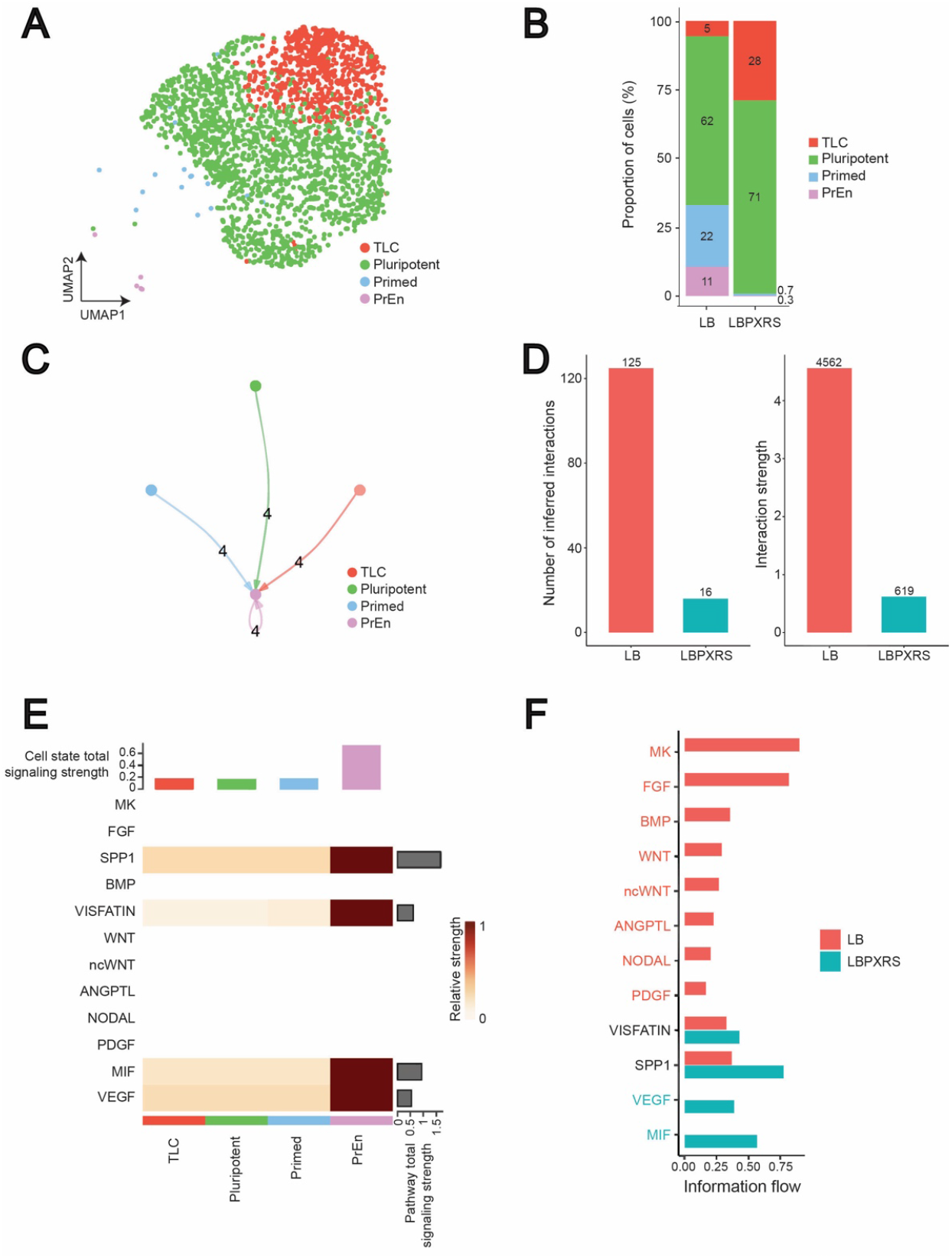
Comparative analysis of the intercellular communication landscape between LB and LBPXRS conditions. **(A)** UMAP visualization of the single-cell RNA-seq data of ESCs from the LBPXRS condition. **(B)** Barplot comparing the proportions of identified cell types in LB and LBPXRS. **(C)** Circle plots depicting the number of inferred intercellular interactions among cell types in the LBPXRS condition as computed by CellChat. Edge thickness corresponds to interaction strength, while the arrow signifies the direction of the communication. **(D)** Quantitative comparison of the total number of interactions and overall communication strength amongst LB and LBPXRS conditions. **(E)** Overview of signalling pathway involvement in LBPXRS. The bar plot on the top shows the total signalling strength of a cell group by summarizing all signalling pathways displayed in the heatmap. The right grey bar plot shows the total signalling strength of a signalling pathway by summarizing all cell groups displayed in the heatmap. **(F)** Stacked barplot of the information flow associated with each pathway across LB and LBPXRS conditions.

Consistent with reduced heterogeneity, CellChat analysis of LBPXRS scRNA-seq data showed a simplified and sparsely connected signalling network, indicating that intercellular communication is significantly restricted **(Fig. 4C)**. Quantitative assessment of global communication metrics further supported this observation, with LB exhibiting substantially more interactions (125 vs. 16) and stronger overall interaction strength (4.562 vs. 0.619) compared to LBPXRS **(Fig. 4D)**. Thus, while LB fosters a dynamic and responsive niche environment that supports greater diversity of cell states, LBPXRS restricts signalling activity and promotes a more uniform cellular landscape.

To further dissect these differences, we performed pathway-level comparative analysis **(Fig. 4E)**. LB cultures engaged a broad array of signalling routes, including FGF, BMP, WNT, NODAL, and PDGF, whereas these pathways were absent under LBPXRS **(Figs. 2C** and **4E-F)**. Instead, exclusive communication in LBPXRS was limited to MIF and VEGF signalling, while common pathways between the two conditions included SPP1 and VISFATIN. Notably, interaction strength appeared inflated for the PrEn state due to its very low cell count, underscoring the importance of accounting for relative cell-type abundance in CCC inference. These findings highlight that the molecular signals supporting cell identity and intercellular communication are strongly shaped by culture conditions, with LB providing a signalling-rich environment that promotes plasticity and cell state transitions, whereas LBPXRS enforces a signalling-restricted space characterized by reduced cell state heterogeneity.

These data indicate that inhibition of FGF, WNT, and NODAL (TGF-β) pathways in LBPXRS culture constrains intercellular communication, leading to a simplified signalling network and restricted usage of pathways. This reduction in communication capacity underlies the observed loss of heterogeneity and emphasizes the critical role of culture-dependent signalling in shaping cellular identity landscapes.

### BMP and NODAL (TGF-β) driven CCC regulates the TLCs

Focusing on the LB data, we next examined ligand-receptor interactions of the BMP, TGF-β, WNT, FGF, and PDGF pathways, using an interaction strength threshold of 0.4, to identify the exclusive and most confident communication routes involving TLCs. We observed limited ligand-receptor interactions, mediated by *Bmp2/4-Hfe2*, with TLCs as senders and pluripotent and primed cells serving as receivers **(Fig. 5A)**. On the contrary, we observed multiple ligand-receptor interactions, involving *F2-Gp5, Nrtn-Ret, Tgfb3-Tgfbr1* and *Tgfb3-Tgfbr2*, with pluripotent, PrEn, and primed cells as senders and TLCs as receivers **(Fig. 5A)**. This suggested a potentially unique incoming signalling axis to TLCs. These patterns suggest that TLCs act as both senders and receivers, engaging in multiple signalling systems to influence the states of their neighbouring cells.

**Figure 5:**
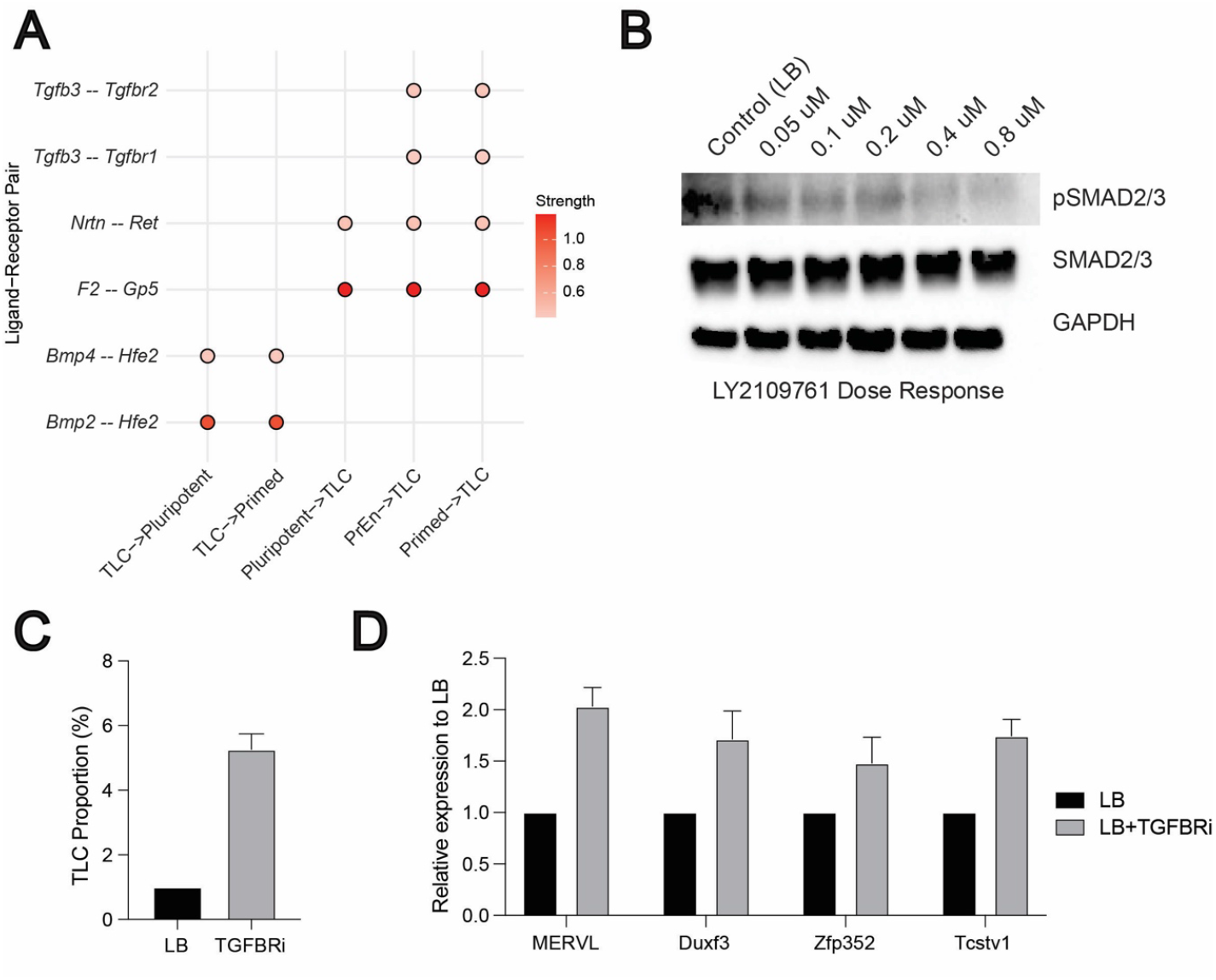
BMP and NODAL (TGF-β) driven CCC regulates the TLCs. **(A)** A compiled Dot plot of differentially expressed ligand–receptor (LR) pairs across “TLC-senders” and “TLC-receivers” from FGF, BMP, WNT, NODAL (TGF-β), and PDGF signalling pathways. Scale represents the strength of the interaction potential. **(B)** Western blot analysis of NODAL (TGF-β) pathways activation in ESCs treated with increasing does of the TGF-β receptor inhibitor LY2109761. **(C)** Bar graph showing the proportion of TLCs (MERVL-tdTomato positive cells) in LB and TGFBRi conditions. **(D)** Relative mRNA levels of TLC genes: *MERVL, Duxf3, Zfp352, and Tcstv1* quantified by qRT-PCR, normalized to housekeeping genes (*Gapdh*), and fold change is relative to LB controls (n=3). In both C and D, error bars represent the standard error of the mean.

We have previously demonstrated the crucial role of BMP2/4 for inducing the TLC state (17). Inhibition of BMP signalling using LDN193189, a selective inhibitor of BMP receptors (35), in the LB condition drastically reduced the TLC proportions. In addition, increasing the dose of BMP correspondingly increased the proportion of TLCs and the expression of TLC genes (17). These experimental data further render validation of the above predicted role of BMP signalling in CCC, consistent with its predicted role in influencing the receiver cell states to promote the TLC state (17). To similarly validate the predicted role of NODAL (TGF-β) signalling, we next targeted the TGFBR1/2 (“receiver” pair) **(Fig. 5A)**. We chose LY2109761, a known specific small molecule inhibitor, to inhibit TGFBR1/2 activation (36). We used ESCs harbouring MERVL-tdTomato live-cell fluorescent reporter to evaluate the changes in TLC proportions using flow cytometry (17, 37). First, by performing a dose-response analysis, we identified 0.8 μM LY2109761 as the concentration that maximally inhibits the activation of NODAL (TGF-β) signalling (**Fig. 5B**). Next, we assessed the change in TLC proportion using flow cytometry and the expression of TLC genes by qPCR upon inhibition of TGF-β signalling with LY2109761. We observed a ∼5-fold increase in TLC proportion and ∼2-fold increase in the expression of multiple TLC genes (*MERVL, Duxf3, Zfp352, Tcstv1*) **(Fig. 5C-D)**. This data suggests that TGF-β-mediated communication to the TLCs acts as a negative regulator of the TLC state.

Together, these results demonstrate that TLC abundance is tightly regulated by the BMP and NODAL (TGF-β) signalling as positive and negative regulators, respectively. It also demonstrates that TLCs integrate multiple extrinsic growth factor signalling in order to either maintain or exit the TLC state.

### In-silico perturbation of BMP-associated transcription factors reveals cell state transition dynamics

Cell signalling often orchestrates cell fate decisions via specific transcription factors (TF). To determine how BMP signalling influences cell fate dynamics of TLCs and other cell states, we utilized CellOracle, which predicts cell fate dynamics by integrating gene regulatory network inference (25). Since CellOracle requires a pseudotime variable as input to construct its developmental vector field, we first inferred pseudotime trajectories with Monocle (26), using TLCs as the source to anchor developmental time. The Monocle graph showed a main bifurcation proximal to the pluripotent compartment, splitting into a branch that remains within the pluripotent space—aligning with a proliferative/plastic subcluster rather than a separate lineage—and another branch moving toward the PrEn state **(Fig. 6A)**. The predicted pseudotime trajectory closely matches the natural developmental cell fate transitions of preimplantation development (17, 38). To establish a dynamics-aware baseline that is unaffected by perturbations, we calculated CellOracle’s digitized developmental flow, a vector field derived directly from the transcriptomic landscape using the Monocle pseudotime variable as input. This baseline flow recapitulated the two main routes seen with Monocle—one confined within the pluripotent niche and another diverging toward PrEn—providing a reference framework to measure the effects of perturbations **(Fig. 6B)**.

**Figure 6:**
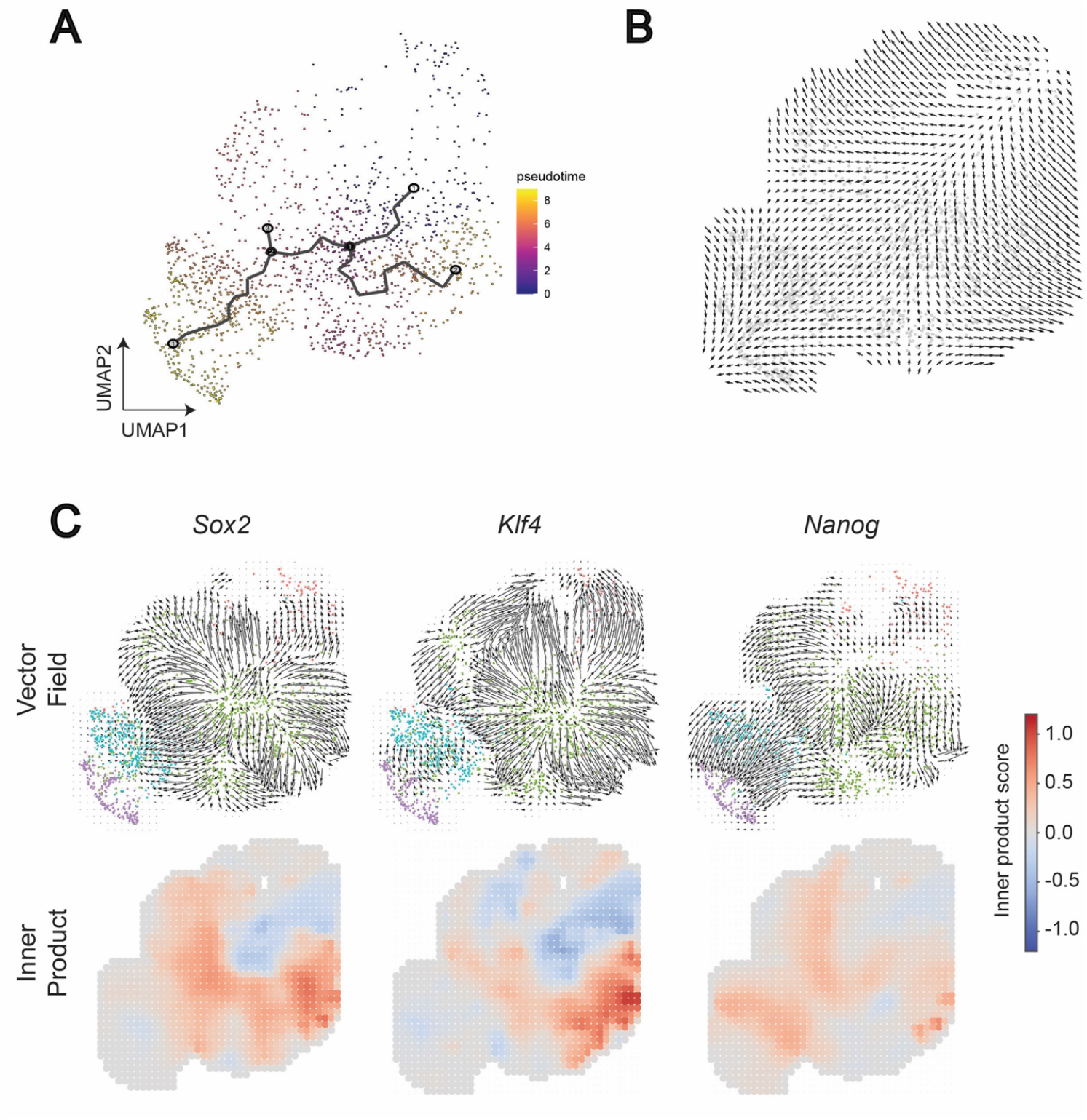
Developmental flow inference and cell state gene perturbation simulations in LB. **(A)** Monocle pseudotime inference. The TLC population was designated as the source state to reconstruct developmental trajectories. **(B)** CellOracle baseline developmental flow. CellOracle computed a pseudotime-like baseline vector field directly from the transcriptome manifold, visualized on a digitized grid. This developmental flow reflects the intrinsic transcriptional dynamics of the system. **(C)** CellOracle transcription factor knockout simulations. For each pluripotent cell state transcription factor (*Sox2, Klf4, Nanog)*, perturbation outcomes are shown in two subpanels: (top) vector fields illustrating local shifts in developmental trajectories compared to baseline flow, (bottom) inner product heatmaps quantifying alignment between perturbation vectors and baseline flow (positive values indicate reinforcement, negative values indicate opposition).

We estimated the point of divergence for each gene of interest (the start of deviation from the developmental flow) and the inner product (positive = reinforcement of the developmental flow; negative = opposition against the developmental flow). To verify that the CellOracle correctly predicts the stem cell state dynamic transitions, we first perturbed the well-known pluripotency TFs. The perturbation vector fields for pluripotent genes *Sox2, Klf4*, and *Nanog* (39-41) all diverge from within the pluripotent population **(Fig. 6C)**. *Sox2* and *Klf4* show broad negative inner products across the pluripotent population, with convergence in the primed state (positive inner product). *Nanog* generally exhibits weaker negative inner products, while positive inner products extend into more differentiated states, and the main convergence shifts toward PrEn. These results confirm that CellOracle correctly predicts the gene function for stem cell state transitions.

Next, we tested the role of key TFs controlled by cell signalling pathways LIF and BMP. Perturbation of the LIF pathway TF *Stat3* (42) diverged the vector field in the primed state **(Fig. 7A)**. Inner products are mostly negative in the primed state, with positive alignment in TLC and PrEn, indicating a modest, context-dependent modulation. In contrast, perturbation of BMP pathways TFs, *Smad1* and *Id1*, had a major impact on the vector fields (43, 44). *Smad1* perturbation diverged the vector field in PrEn and showed predominantly negative inner products that converged toward the pluripotent state. *Id1*, another BMP pathway target, diverged in primed and produced strong, extensive negative inner products across primed and pluripotent regions, converging toward the TLC state.

**Figure 7:**
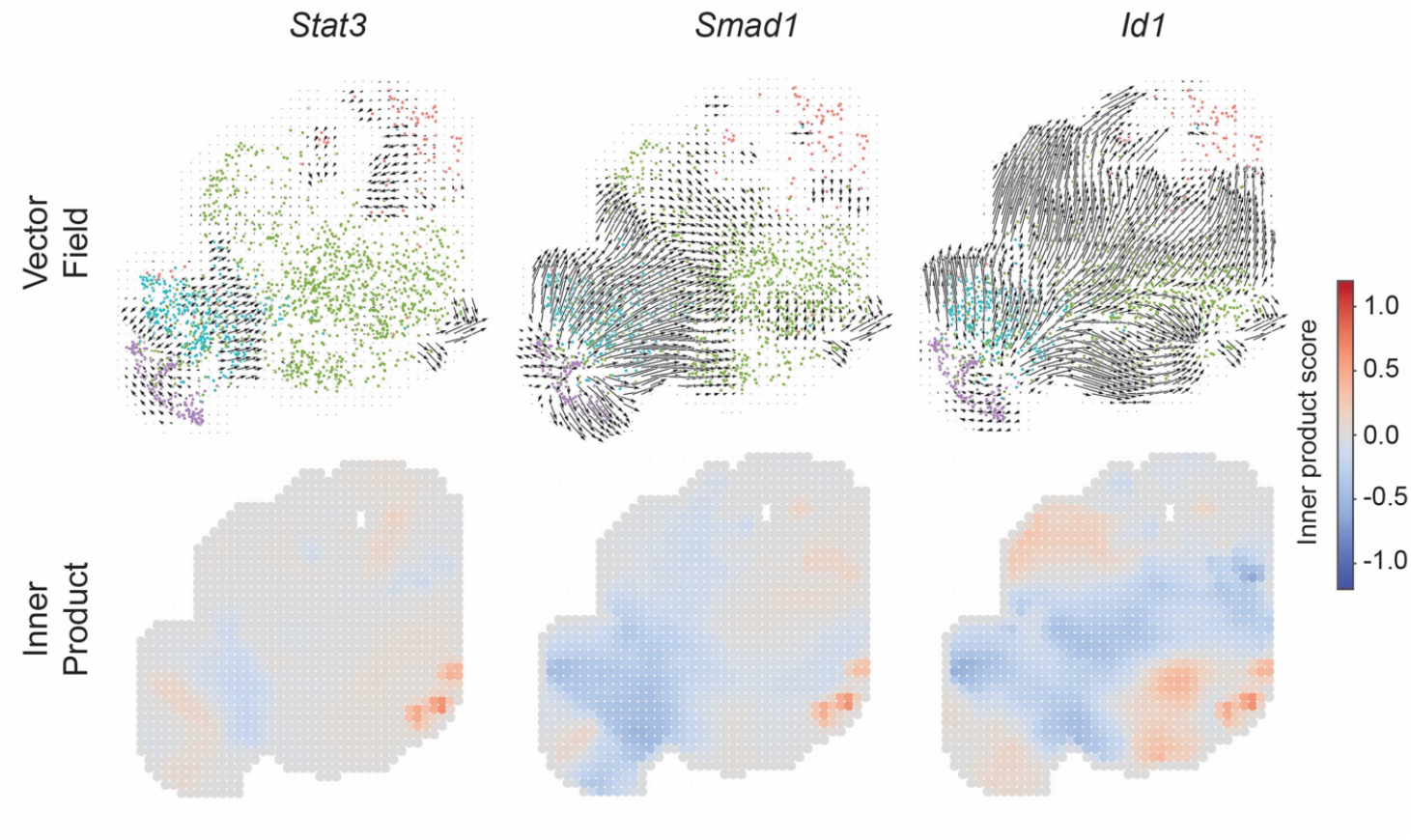
Signalling pathway target transcription factor perturbation simulations in LB. CellOracle transcription factor knockout simulations. For each transcription factor (*Stat3, Smad1* and *Id1*), perturbation outcomes are shown in two subpanels: (top) vector fields illustrating local shifts in developmental trajectories compared to baseline flow, (bottom) inner product heatmaps quantifying alignment between perturbation vectors and baseline flow (positive values indicate reinforcement, negative values indicate opposition).

Together, these data further confirm the crucial role of BMP signalling in regulating the TLC state as well as other stem cell states of early embryonic development (17, 45). They provide a mechanistic model to link normal developmental flow with variable transcriptional responses induced by cell signalling.

## 5) DISCUSSION

Our findings highlight how paracrine signalling influences ESC fate dynamics, with TLCs being continually shaped by signals from their pluripotent, primed, and PrEn neighbours. Such intercellular communication plays a crucial role in early embryonic development. For instance, in the preimplantation embryo, FGF4 secreted by epiblast precursors instructs adjacent inner cell mass cells to adopt a PrEn identity, exemplifying a paracrine-driven lineage specification (46). At the peri-implantation stage, reciprocal signalling between tissues orchestrates morphogenetic remodelling – epiblast-derived factors help sustain extraembryonic lineages, such as trophectoderm and endoderm (47), and emerging morphogen gradients break symmetry to establish body axes (48). Notably, localized BMP4 signals trigger the WNT and NODAL pathways across the epiblast and visceral endoderm, initiating the formation of the anterior-posterior axis (48). Such parallels suggest that spatially confined microenvironments act as information-processing hubs: cells integrate multiple concurrent morphogen inputs, and developmental systems filter out stochastic noise to ensure robust patterning (5). In essence, the embryo’s signalling niches dynamically balance stem cell self-renewal while maintaining developmental equilibrium. This perspective also implies that by manipulating paracrine signalling, we can steer cell plasticity in culture. Indeed, experimentally tuning key pathways (e.g. inhibiting FGF, WNT, and NODAL while enhancing BMP signals) biases ESC populations toward the TLC state (17). Harnessing these principles could enable engineered control of cell fate in vitro, allowing us to recapitulate in vivo cell fate decision-making of embryonic development.

In our study, we observed that increased cellular heterogeneity is associated with a more diverse intercellular communication network. This likely reflects the contribution of distinct transcriptional profiles across individual cells, which together enhance the stability and resilience of the signalling landscape. Previous work has highlighted the role of paracrine and autocrine signalling in regulating pluripotency and lineage commitment (12, 49), and our findings extend this by suggesting that heterogeneity itself may serve as a stabilizing factor in maintaining pluripotent states. The importance of this principle becomes evident in disease contexts such as intrahepatic cholangiocarcinoma (ICC), where reduced cellular diversity coincides with a less intricate outgoing signalling network (50). Interestingly, the cancer cell populations exhibited greater heterogeneity and stronger outgoing signalling compared to their normal counterparts, suggesting that heterogeneity may confer adaptive advantages in pathological settings. Collectively, these results support the notion that cellular heterogeneity functions as a general mechanism for population stability and adaptability. This has important implications not only for understanding tumour progression but also for optimizing stem cell culture conditions and developing therapeutic strategies that seek to preserve or manipulate cell states through modulation of intercellular communication.

Our findings reveal distinct effects of NODAL (TGF-β) pathway inhibition on TLC regulation, underscoring their differential contributions to cell state control. Inhibition of TGF-β signalling produced both a marked increase in TLC frequency and an upregulation of TLC gene expression, indicating that suppression of this pathway promotes the expansion or stabilization of the TLC population. These findings suggest that TGF-β signalling negatively regulates both the transcriptional and cellular maintenance of TLCs. Together, the data highlight a potential inhibitory role of TGF-β signalling in TLC maintenance.

The in-silico gene perturbation behaviours we observed between cell state genes and pathway targets can be explained by their distinct roles in regulation. Perturbations of pluripotent cell state markers were first performed to validate that the model correctly reproduces expected transcriptional behaviours within the pluripotent population, ensuring that predicted cell fate shifts reflect biologically coherent responses. Cell state genes, such as *Sox2, Klf4*, and *Nanog*, primarily serve as indicators of pluripotent identity. As expected, their perturbation triggers a pluripotency exit, resulting in positive inner products that align with differentiation. This finding also aligns with previous research that suggests *Nanog* is not strictly essential for self-renewal, as altering its levels can influence lineage commitment((51), and that *Sox2* and *Klf4* primarily determine pluripotent identity((52). We then extended the perturbations to pathway targets like *Stat3, Smad1*, and *Id1*, which are TFs that actively help stabilize or redirect cell fate decisions. For instance, *Stat3* activation downstream of LIF is both necessary and sufficient to maintain self-renewal((42, 53), and BMP signalling maintains pluripotency through *Smad1* and *Id1*-mediated suppression of stem cell differentiation((44). These results imply that perturbing specific signalling pathway TFs dynamically alter the natural developmental cell fate transitions.

### Perspectives and Limitations

While this study provides insights into intercellular communication in ESCs, several limitations should be considered. First, our analysis does not address the effects of autocrine signalling on the TLC state. Autocrine loops are known to influence cell fate decisions(12), and incorporating this dimension would refine our understanding of how individual cells integrate self-derived vs extrinsic cues. Second, all our validations were performed in a two-dimensional *in vitro* setting. Although 2D cultures are tractable and widely used, they lack the spatial architecture and mechanical context of early embryonic environments. Extending these approaches to three-dimensional models such as embryoid bodies or organoid systems would provide a more physiologically relevant framework and enhance the translational applicability of our findings.

## REFERENCES

1. Casey MJ, Stumpf PS, MacArthur BD. Theory of cell fate. Wiley Interdiscip Rev Syst Biol Med. 2020;12(2):e1471.

2. Wu S, Schmitz U. Single-cell and long-read sequencing to enhance modelling of splicing and cell-fate determination. Comput Struct Biotechnol J. 2023;21:2373–80.

3. Lee J, Kim N, Cho KH. Decoding the principle of cell-fate determination for its reverse control. NPJ Syst Biol Appl. 2024;10(1):47.

4. Haghverdi L, Ludwig LS. Single-cell multi-omics and lineage tracing to dissect cell fate decision-making. Stem Cell Reports. 2023;18(1):13–25.

5. Davies AE, Albeck JG. Microenvironmental Signals and Biochemical Information Processing: Cooperative Determinants of Intratumoral Plasticity and Heterogeneity. Front Cell Dev Biol. 2018;6:44.

6. Su J, Song Y, Zhu Z, Huang X, Fan J, Qiao J, et al. Cell-cell communication: new insights and clinical implications. Signal Transduct Target Ther. 2024;9(1):196.

7. Singer SJ. Intercellular communication and cell-cell adhesion. Science. 1992;255(5052):1671–7.

8. Wu G, Liang Y, Xi Q, Zuo Y. New Insights and Implications of Cell-Cell Interactions in Developmental Biology. Int J Mol Sci. 2025;26(9).

9. Cooper GM. The Cell: A Molecular Approach. 2nd Edition ed 2000.

10. Movasat H, Giacopino E, Shahdoost A, Dorri Nokoorani Y, Abrbekouh AH, Tahamtani Y, et al. A systems view of cellular heterogeneity: Unlocking the “wheel of fate”. Cell Syst. 2025;16(6):101300.

11. Morrison SJ, Spradling AC. Stem cells and niches: mechanisms that promote stem cell maintenance throughout life. Cell. 2008;132(4):598–611.

12. Przybyla L, Voldman J. Probing embryonic stem cell autocrine and paracrine signaling using microfluidics. Annu Rev Anal Chem (Palo Alto Calif). 2012;5:293–315.

13. (US) NRC, Research IoMUCotBaBAoSC. Stem cells and the future of regenerative medicine. Washington, D.C: National Academy Press; 2002.

14. Liang G, Zhang Y. Embryonic stem cell and induced pluripotent stem cell: an epigenetic perspective. Cell Res. 2013;23(1):49–69.

15. Romito A, Cobellis G. Pluripotent Stem Cells: Current Understanding and Future Directions. Stem Cells Int. 2016;2016:9451492.

16. Lanner F, Rossant J. The role of FGF/Erk signaling in pluripotent cells. Development. 2010;137(20):3351–60.

17. Meharwade T, Joumier L, Parisotto M, Huynh V, Lummertz da Rocha E, Malleshaiah M. Cross-activation of FGF, NODAL, and WNT pathways constrains BMP-signaling-mediated induction of the totipotent state in mouse embryonic stem cells. Cell Rep. 2023;42(5):112438.

18. Singer ZS, Yong J, Tischler J, Hackett JA, Altinok A, Surani MA, et al. Dynamic heterogeneity and DNA methylation in embryonic stem cells. Mol Cell. 2014;55(2):319–31.

19. Chehelgerdi M, Behdarvand Dehkordi F, Chehelgerdi M, Kabiri H, Salehian-Dehkordi H, Abdolvand M, et al. Exploring the promising potential of induced pluripotent stem cells in cancer research and therapy. Mol Cancer. 2023;22(1):189.

20. Yang S, Cho Y, Jang J. Single cell heterogeneity in human pluripotent stem cells. BMB Rep. 2021;54(10):505–15.

21. Lu F, Zhang Y. Cell totipotency: molecular features, induction, and maintenance. Natl Sci Rev. 2015;2(2):217–25.

22. Jovic D, Liang X, Zeng H, Lin L, Xu F, Luo Y. Single-cell RNA sequencing technologies and applications: A brief overview. Clin Transl Med. 2022;12(3):e694.

23. Jin S, Guerrero-Juarez CF, Zhang L, Chang I, Ramos R, Kuan CH, et al. Inference and analysis of cell-cell communication using CellChat. Nat Commun. 2021;12(1):1088.

24. Browaeys R, Saelens W, Saeys Y. NicheNet: modeling intercellular communication by linking ligands to target genes. Nat Methods. 2020;17(2):159–62.

25. Kamimoto K, Stringa B, Hoffmann CM, Jindal K, Solnica-Krezel L, Morris SA. Dissecting cell identity via network inference and in silico gene perturbation. Nature. 2023;614(7949):742–51.

26. Trapnell C, Cacchiarelli D, Grimsby J, Pokharel P, Li S, Morse M, et al. The dynamics and regulators of cell fate decisions are revealed by pseudotemporal ordering of single cells. Nat Biotechnol. 2014;32(4):381–6.

27. Huang G, Ye S, Zhou X, Liu D, Ying QL. Molecular basis of embryonic stem cell self-renewal: from signaling pathways to pluripotency network. Cell Mol Life Sci. 2015;72(9):1741–57.

28. Lafyatis R. Transforming growth factor β—at the centre of systemic sclerosis. Nature Reviews Rheumatology. 2014;10(12):706–19.

29. Xue C, Chu Q, Shi Q, Zeng Y, Lu J, Li L. Wnt signaling pathways in biology and disease: mechanisms and therapeutic advances. Signal Transduction and Targeted Therapy. 2025;10(1):106.

30. Ornitz DM, Itoh N. The Fibroblast Growth Factor signaling pathway. Wiley Interdiscip Rev Dev Biol. 2015;4(3):215–66.

31. Xu H, Liang H. The regulation of totipotency transcription: Perspective from in vitro and in vivo totipotency. Front Cell Dev Biol. 2022;10:1024093.

32. Chen G, Yin S, Zeng H, Li H, Wan X. Regulation of Embryonic Stem Cell Self-Renewal. Life (Basel). 2022;12(8).

33. Karaulanov EE, Böttcher RT, Niehrs C. A role for fibronectin-leucine-rich transmembrane cell-surface proteins in homotypic cell adhesion. EMBO Rep. 2006;7(3):283–90.

34. Yang R, Chen Y, Chen D. Biological functions and role of CCN1/Cyr61 in embryogenesis and tumorigenesis in the female reproductive system (Review). Mol Med Rep. 2018;17(1):3–10.

35. Cuny GD, Yu PB, Laha JK, Xing X, Liu J-F, Lai CS, et al. Structure–activity relationship study of bone morphogenetic protein (BMP) signaling inhibitors. Bioorganic & Medicinal Chemistry Letters. 2008;18(15):4388–92.

36. Melisi D, Ishiyama S, Sclabas GM, Fleming JB, Xia Q, Tortora G, et al. LY2109761, a novel transforming growth factor beta receptor type I and type II dual inhibitor, as a therapeutic approach to suppressing pancreatic cancer metastasis. Mol Cancer Ther. 2008;7(4):829–40.

37. Macfarlan TS, Gifford WD, Driscoll S, Lettieri K, Rowe HM, Bonanomi D, et al. Embryonic stem cell potency fluctuates with endogenous retrovirus activity. Nature. 2012;487(7405):57–63.

38. Du P, Wu J. Hallmarks of totipotent and pluripotent stem cell states. Cell Stem Cell. 2024;31(3):312–33.

39. Avilion AA, Nicolis SK, Pevny LH, Perez L, Vivian N, Lovell-Badge R. Multipotent cell lineages in early mouse development depend on SOX2 function. Genes Dev. 2003;17(1):126–40.

40. Di Giammartino DC, Kloetgen A, Polyzos A, Liu Y, Kim D, Murphy D, et al. KLF4 is involved in the organization and regulation of pluripotency-associated three-dimensional enhancer networks. Nat Cell Biol. 2019;21(10):1179–90.

41. Pan G, Thomson JA. Nanog and transcriptional networks in embryonic stem cell pluripotency. Cell Res. 2007;17(1):42–9.

42. Niwa H, Burdon T, Chambers I, Smith A. Self-renewal of pluripotent embryonic stem cells is mediated via activation of STAT3. Genes Dev. 1998;12(13):2048–60.

43. Trompouki E, Bowman Teresa V, Lawton Lee N, Fan Zi P, Wu D-C, DiBiase A, et al. Lineage Regulators Direct BMP and Wnt Pathways to Cell-Specific Programs during Differentiation and Regeneration. Cell. 2011;147(3):577–89.

44. Ying QL, Nichols J, Chambers I, Smith A. BMP induction of Id proteins suppresses differentiation and sustains embryonic stem cell self-renewal in collaboration with STAT3. Cell. 2003;115(3):281–92.

45. Graham SJL, Wicher KB, Jedrusik A, Guo G, Herath W, Robson P, et al. BMP signalling regulates the pre-implantation development of extra-embryonic cell lineages in the mouse embryo. Nature Communications. 2014;5(1):5667.

46. Kang M, Garg V, Hadjantonakis AK. Lineage Establishment and Progression within the Inner Cell Mass of the Mouse Blastocyst Requires FGFR1 and FGFR2. Dev Cell. 2017;41(5):496–510.e5.

47. Vrij EJ, Scholte Op Reimer YS, Roa Fuentes L, Misteli Guerreiro I, Holzmann V, Frias Aldeguer J, et al. A pendulum of induction between the epiblast and extra-embryonic endoderm supports post-implantation progression. Development. 2022;149(20).

48. Cheng D, Clark CT, Smith Q. Advances in engineered models of peri-gastrulation. iScience. 2025;28(6):112659.

49. Dalton S. Signaling networks in human pluripotent stem cells. Curr Opin Cell Biol. 2013;25(2):241–6.

50. Zhou ZQ, Zhang Y, Xu ZY, Tang XL, Chen XH, Guan J, et al. Dissecting cellular heterogeneity and intercellular communication in cholangiocarcinoma: implications for individualized therapeutic strategies. Front Genet. 2023;14:1241834.

51. Chambers I, Tomlinson SR. The transcriptional foundation of pluripotency. Development. 2009;136(14):2311–22.

52. Takahashi K, Yamanaka S. Induction of Pluripotent Stem Cells from Mouse Embryonic and Adult Fibroblast Cultures by Defined Factors. Cell. 2006;126(4):663–76.

53. Matsuda T, Nakamura T, Nakao K, Arai T, Katsuki M, Heike T, et al. STAT3 activation is sufficient to maintain an undifferentiated state of mouse embryonic stem cells. The EMBO Journal. 1999;18(15):4261–9-9.

